# A learnable transition from low temperature to high temperature proteins with neural machine translation

**DOI:** 10.1101/2024.02.06.579188

**Authors:** Evan Komp, Christian Phillips, Humood N. Alanzi, Marlo Zorman, David A. C. Beck

## Abstract

This work presents Neural Optimization for Melting-temperature Enabled by Leveraging Translation (NOMELT), a novel approach for designing and ranking high-temperature stable proteins using neural machine translation. The model, trained on over 4 million protein homologous pairs from organisms adapted to different temperatures, demonstrates promising capability in targeting thermal stability. A designed variant of the *Drosophila melanogaster* Engrailed Homeodomain shows increased stability at high temperatures, as validated by estimators and molecular dynamics simulations. Furthermore, NOMELT achieves zero-shot predictive capabilities in ranking experimental melting and half-activation temperatures across two protein families. It achieves this without requiring extensive homology data or massive training datasets as do existing zero-shot predictors by specifically learning thermophilicity, as opposed to all natural variation. These findings underscore the potential of leveraging organismal growth temperatures in context-dependent design of proteins for enhanced thermal stability.

## Introduction

Proteins have been leveraged in human society for a variety of applications such as therapeutics, food processing, textiles, commodity materials, waste remediation, and have the potential for future impacts such as plastic upcycling.[1–8] The efficacy of these applications is often hampered by a key challenge: enhancing the high temperature thermal stability of active proteins towards more favorable system conditions. Researchers have been searching for methods to rapidly increase the thermal stability of proteins for decades, with many contradictory findings, revealing that there are no generally applicable “rules” that can predictably improve thermal stability. Instead, the thermal stability of each protein is convoluted and dependent on the protein itself, i.e. each protein is a unique engineering challenge.[9–15]

Recently, deep learning has been an accelerating force for protein engineers, and large attention based models are at the forefront.[16–18] Successes include supervised predictors of a property of interest, including thermal stability,[19,20] zero-shot predictors of the same,[21,22] structure prediction models,[23,24] and sequence design models.[25,26] Existing supervised strategies help rank proteins among a pool of variants after training on a specific thermal stability target, but require time and resource intensive labeled data from the specific protein of interest to be accurate.[27–29] Zero-shot predictors remove the need for labeled data by either learning from evolutionary scale sequence datasets or by conditioning on homologs of the protein of interest and impressively achieve some predictive performance on observable properties. Unfortunately, the latter requires many known homologs to retain fidelity, and both are not strictly targeting thermal stability, unable to reliably rank the global optimum among a dataset.[27] Structure to sequence designers are able to create amino acid sequences that likely fold to a desired 3D structure at ambient temperatures, but do not specifically target high temperature stability and none have yet been show to reliably retain function at high temperatures to the best of our knowledge. These deficiencies motivate the current work - can evolutionary sequence data be organized in such a way that we can accelerate the design of high temperature proteins in a context dependent manner without supervised training?

Presented is a method for designing and ranking high temperature proteins by learning a translation between ambient and high temperature protein space using neural machine translation: Neural Optimization for Melting-temperature Enabled by Leveraging Translation (NOMELT). By training a neural machine translator[30] on a dataset greater than 4 million pairs of protein homologs, where one protein occurs in an ambient temperature prokaryote and the other in a high temperature prokaryote, we show that a protein language model can target thermal stability. The model was used to autoregressively inject mutations to create a variant of *Drosophila melanogaster* Engrailed Homeodomain (EnHD) that is stable at a higher temperature according to both qualitative estimators and molecular dynamics simulations. It is also shown to have zero-shot predictive capabilities for ranking variants by experimental melting temperatures and catalytic half activation temperatures for two different protein families. Unlike existing zero-shot predictors, NOMELT does not require homologs as input and was not trained on tens of millions data points. These results suggest that leveraging organism growth temperature improves the richness of data for designing high temperature proteins.

## Results and Discussion

### Thermophilic sequences can be recapitulated from mesophilic homologs

The NOMELT model was trained to, given an input mesophilic protein sequence, construct a thermophilic homolog by causally building the amino acids N to C terminus. It is known that proteins among a family can have as little as 20% identity and still retain same-or-similar function, and that residues on the length of the protein can occupy a distribution of amino acids and can include gaps and insertions between sequences.[31] Thus, we would not expect nor want the model to perfectly recover every thermophilic amino acid in a particular protein pair. Instead, the data that the model was trained on is redundant among protein pairs, as seen in Figure 1, where the probability density of the number of times a particular sequence is seen in the training set for mesophilic and thermophilic sequences is shown. Each thermophilic sequence to be constructed has on average 103.6 mesophilic counterparts, and 7.1 for the inverse. This is desirable as NOMELT has some redundancy across evolutionary space to learn from.

**Figure 1:**
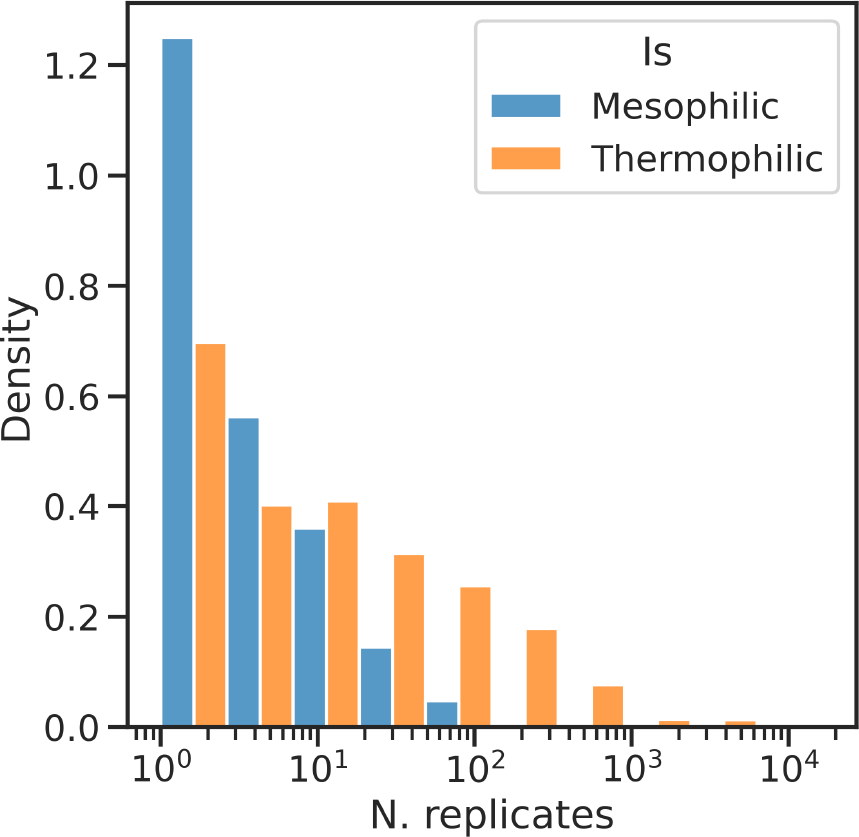
Density of replicate counts for mesophilic and thermophilic sequences occurring in the training set. The most common number of replicates for both the input and the target sequences is one, however the majority of proteins occur multiple times (paired to a different homolog) in the training set. Thermophilic replicates are right skewed, as homologous protein pairs were identified for a much larger group of mesophilic sequences to a smaller set of thermophilic ones.

To evaluate the model’s ability to recapitulate thermophilic counterparts, we consider a held out test set of protein clusters. The test set was extracted from the overall dataset by 50% identity clusters such that similar sequences were not observed across development splits. Further, to reduce evaluation bias towards highly represented protein families, only a single sequence from each cluster was selected, see Methods for details. On this test set we compute a number of metrics by comparing the thermophilic sequence to the model’s output, see Table 1. The test loss is determined causally with teacher forcing for previous amino acids, while all other metrics are determined by comparing to a BEAM search over autoregressive sequence translations, yielding a single translated sequence.[32]

**Table 1.** Scores of the model for recapitulating the test set. “Residue” scores are computed on a per-amino acid basis, “Sequence” are computed by comparing full sequences, and “Structure” are derived from comparison of ESMFold structure prediction. *Does not use the NOMELT model, but instead natural variation over thermophilic homologs.

| Test Metric | Value | Type |
| --- | --- | --- |
| Cross Entropy Loss | 0.90 | Residue |
| MSA Cross Entropy Loss | 0.94 | *Natural\**, Residue |
| Transcription Error Rate | 47% | Sequence |
| Sequence Identity | 43% | Sequence |
| Bits per residue, BLOSUM62[36] | 2.4 | Sequence |
| Jenson-Shannon Secondary Structure | 0.01 | Structure |
| FATCAT Structural Alignment P-value | 0.01 | Structure |

The model generates sequences with a 2.3% difference in length (shorter or longer) compared to the true thermophilic sequence, and has a transcription error rate of 47%, indicating that 53% of the time the model places the exactly correct amino acid in the correct position. Note that the error rate does not consider homology/analogous amino acids, or gaps and deletions, and is normalized to the total number of residues in the test set as opposed to the number of sequences. In order to ground the model’s understanding of the meso to thermo translation space, we leverage known natural variation to qualitatively compare to the test loss value. An MSA was built for each thermophilic sequence by searching the entire thermophilic dataset of homologs using jackhmmer.[33] The cross entropy for the thermophilic targets over the natural variation, residue-wise, is 0.94 on average. The model’s test categorical cross entropy loss is 0.90, indicating that the model is slightly better at predicting the true amino acid at a position than treating each position independently and sampling the most probable residue from natural amino acid distribution over thermophilic homologs. Critically, the model does not require an MSA of other thermophilic homologs, but a single input mesophilic sequence.

To consider the viability of the model to produce full sequences, as opposed to residue-wise prediction, we also took each generated sequence and aligned it to its true test thermophilic sequence and found full length alignment identities between 4.5% and 100% with an average of 43%, and bits-per-residue of 2.4 according to the BLOSUM62 scoring matrix. Only 19% of the time is the generated sequence further from both the mesophilic and thermophilic sequence in identity than they are from each other. Further we labeled the secondary structure using pyDSSP following ESMFold structure for each generated and thermophilic sequence.[24,34] The difference in distribution over Helix, Strand, and Loop structures for each pair of generated-ground truth sequence is measured by the Jenson-Shannon divergence. With a value of 0.01 on average, the model is correctly transcribing the expected secondary structures in target proteins. Lastly, 3D alignment of the generated and thermophilic predicted structures using FATCAT produces a mean P-value of 0.01 and a max of 0.1, indicating that the generated sequence is close in structure to the ground truth, according to ESMFold and FATCAT.[35]

### Known thermophilic protein attributes are captured by the model

The community has been searching through the pool of thermophilic proteins for years in order to determine the rules and modes of high temperature stability. While the general consensus is that there is no universal set of rules, with high temperature stability being highly sequence-dependent, a few notable observations are widely accepted.[37,38] Firstly, it is known that thermophiles leverage a different distribution of amino acids than do mesophiles.[39] We measured the distributions of amino acids for model-generated sequences and found that the model uses a similar shift in amino acid propensities as has been previously reported most of the time. In Figure 2, we can see the change in AA prevalence between thermophilic and mesophilic for model-generated sequences follows the previously reported values in direction and magnitude. Note that the literature values are derived from only 16 proteomes, and our data differs in magnitude for some amino acids. Model generated sequences follow a similar distribution as our test data.

**Figure 2:**
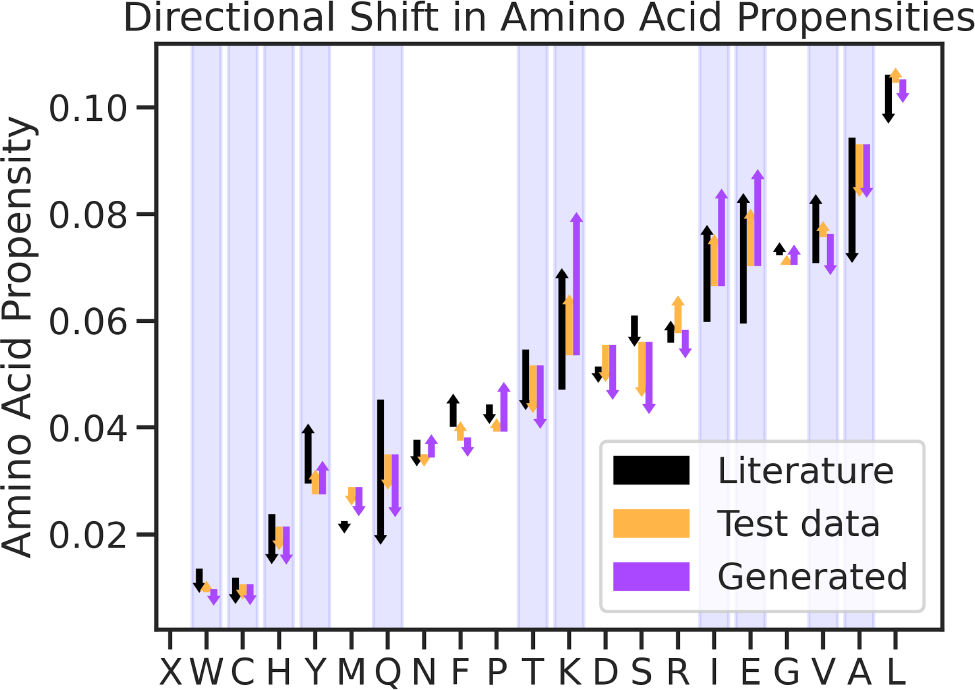
Change in Amino Acids frequencies between mesophilic proteins and thermophilic ones. In black, values derived from 16 proteomes in literature. In orange, the test set of our data, and in purple, generated sequences by our model. Statistically significant shifts determined in literature are highlighted in blue. For almost all significant amino acids, our data has about the same direction in shift as identified from reference proteomes, with most being the correct order of magnitude. The generated sequences recapitulate the distribution observed in our data.

Next, it is accepted that thermophiles tend to leverage disulfide bonds to a greater extent than do mesophiles.[40] The model outputs were probed for the predicted likelihood (Eq. 1) of cysteine to determine if it understands the importance of such bonds for high temperature stability. It was found that the model is significantly more likely to place a cysteine in the sequence to complete a disulfide bond than to place one that does not form a bond. Note that disulfide bonds are posited using the heuristic of 7.5Å alpha carbon distance on the ESMFold predicted structure.[41]

Strictly speaking, the model is trained to convert proteins to look more like thermophilic variants, however we know that these must be high temperature stable, in order for the host organism to thrive. To confirm this, and evaluate the model’s ability to capture it, we adapt the recent mAF-min method by normalizing by sequence length.[42] For the test set examples, we compared the estimated stability of the ground truth thermophilic homolog and the model translation to the mesophilic protein, finding that both are estimated to be more stable with statistical significance. This indicates that the model is successful at learning aspects of stability from thermophilic proteins and applying them to new and diverse families, according to the mAF-min. See Figure 4, below.

**Figure 3:**
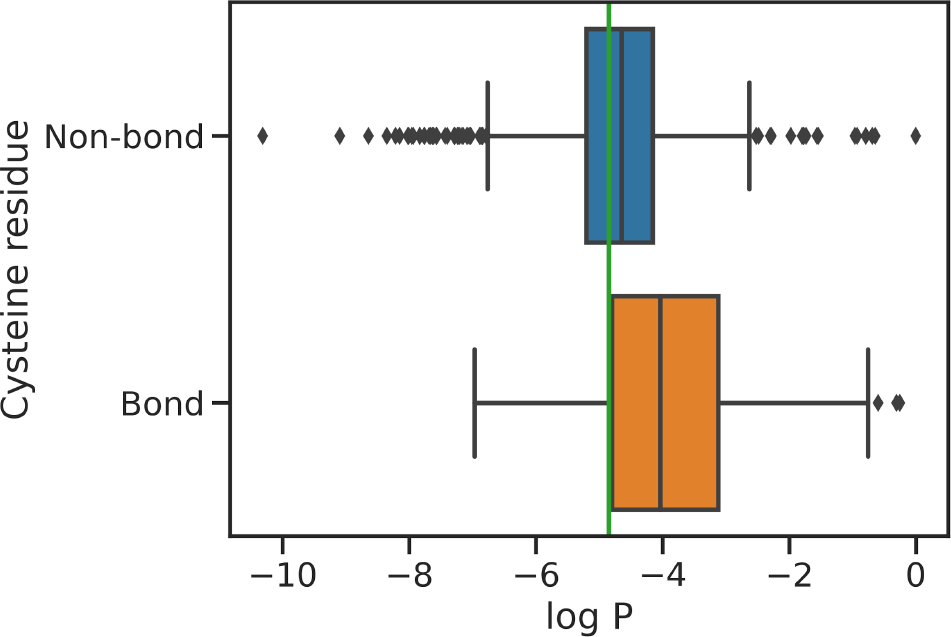
Model predicted log likelihood of Cysteine residues for positions that would or wouldn’t form a disulfide bond with another cysteine in the predicted structure (C_α_ distance <7.5Å[41]). In green, the log likelihood of an amino acid assuming random uniform. For residues where the Cysteine would not form a bond with an existing Cysteine, the model has little Cysteine bias, sometimes predicting Cysteine with high probability and other times choosing different amino acids. For residue positions that would form a disulfide bond, the model has a very heavy bias towards predicting Cysteine.

**Figure 4:**
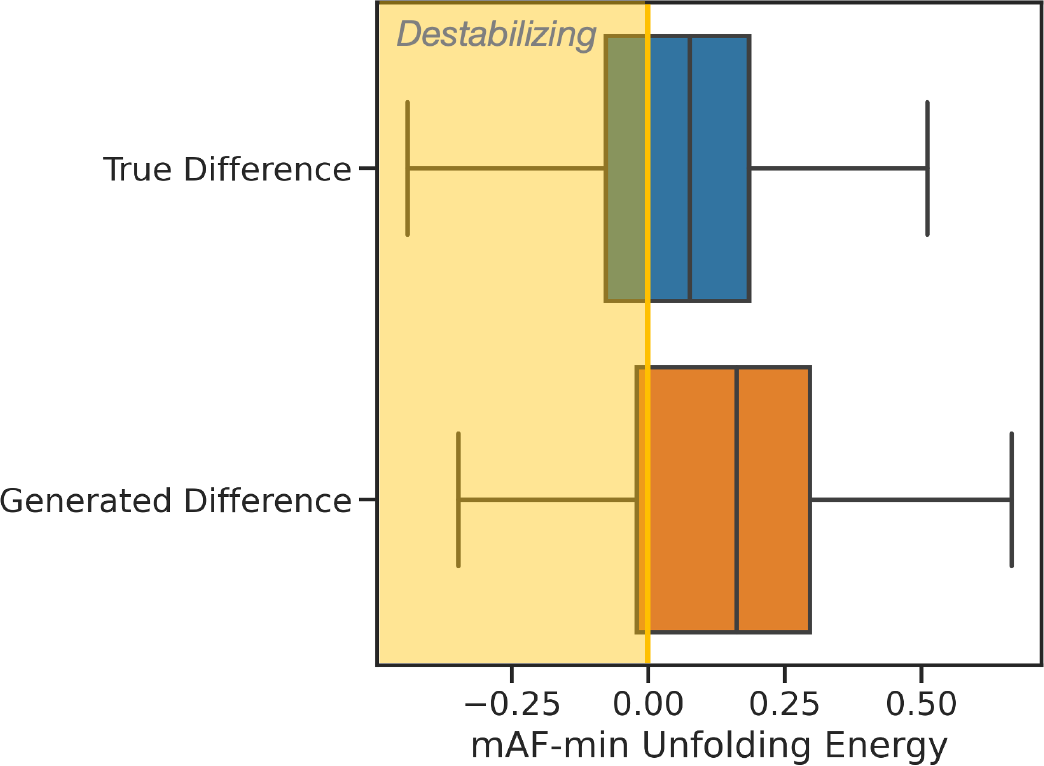
Shift in stability from mesophilic protein, according to the mAF-min method. Ground truth thermophilic sequences, are stabilizing on average, with 56% of examples in the dataset having >95% confidence of having a high folding free energy change. The sequences generated by the model on the test set capture a similar conference of increased stability, with 72% meeting the >95% interval.

#### Case Study 1: Use NOMELT to engineer thermally stabilizing sequence variation

We ran the model on Engrailed homeodomain (En-HD), a three helix bundle transcription factor found in *Drosophila melanogaster*.[43,44] Note that NOMELT’s data contains no eukaryotic proteins, and no sequence in the training set has a BLASTp E-value <1.5 to En-HD.[45] A BEAM search produced a sequence with 14 suggested mutations (insertions and deletions included) when aligned with the wild type protein, as seen in Figure S1. We found that the raw model output was about the same stability as the wild type according to mAF-min with 2.37 and 2.36 respectively (more positive is more stable). For reference, a previously engineered En-HD variant designed by consensus over many homologs has a score of 2.56.[46] While the raw model output did not confer a significant increase in estimated stability, we instead view NOMELT as a generator of variation rather than a single-pass engineer, given that the model may be introducing variation picked up from nature that does not directly impact thermal stability. This yields a library size of 16,384 considering 14 binary mutations to search over.

We conducted 10 rounds of NSGA-ii evolutionary optimization each with a population size of 10 using the mAF-min method as an objective function. These rounds explore only 0.6% of the library, but are able to improve the stability score of En-HD by 71% of the previously engineered variant to 2.51 using NOMELT suggestions. Note that NOMELT requires only a single input sequence, unlike the consensus variant. To ground this result, we repeat the process considering 14 random mutations on the protein, and are unable to improve the stability over the wild type with statistical significance. See Figure 5.

**Figure 5:**
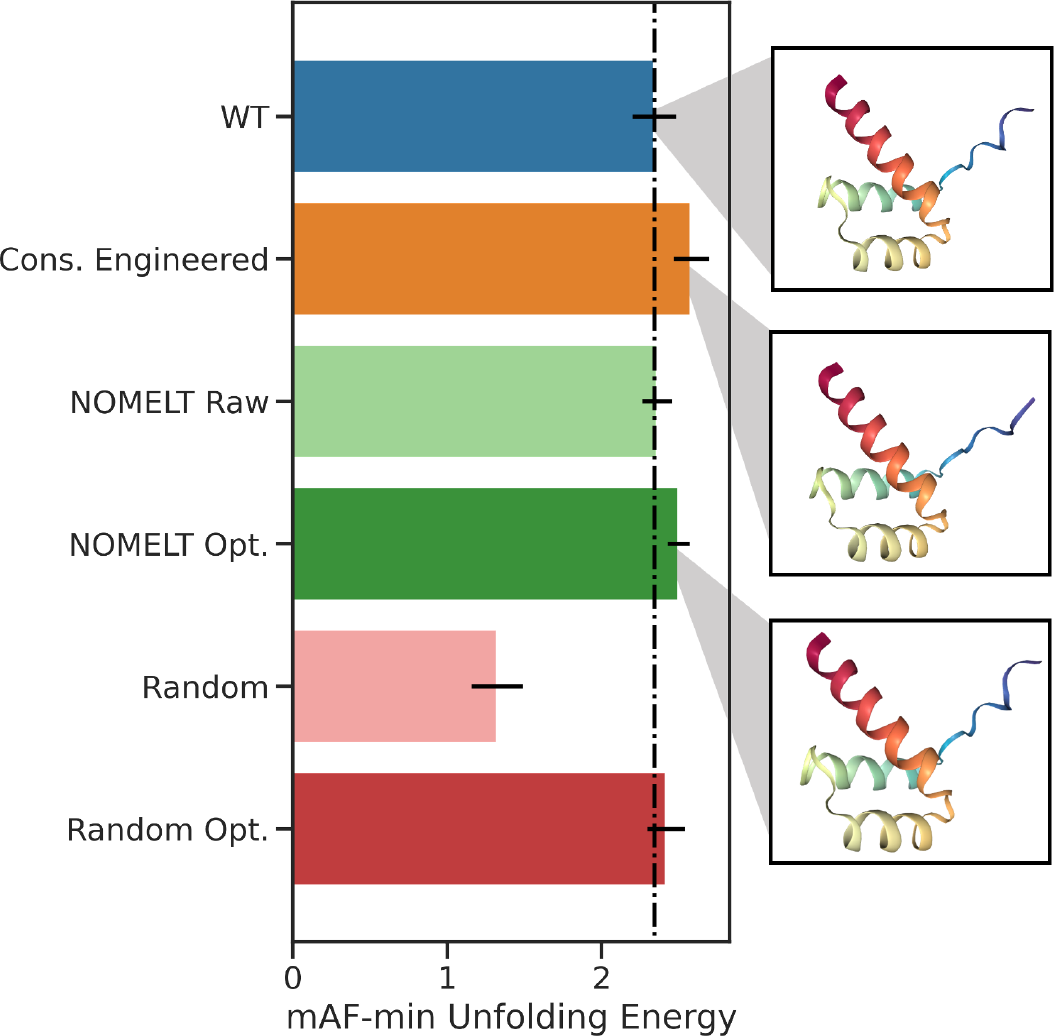
Comparison of stability of various designs according to the mAF-min method’s unfolding free energy change. Error bars indicate 95% CI over the ensemble of AlphaFold structures. Vertical black line is the wild type’s score. The previously engineered variant, by consensus over many homologs (orange) is more stable according to the estimator (P-value 1.7e^-19^). The raw output of the NOMELT model given a single input wild type sequence (light green), which includes 14 mutations involving some insertions, is not stabilizing or destabilizing. After only 100 examples from the search space of 14^2^ mutation permutations have been explored using NSGA-ii (dark green), we achieve a statistically more stable variant than WT (P-value 9.5e^-14^). Introducing the same number of mutations randomly is extremely destabilizing (light red), and running the same number of exploration steps over the random search space cannot improve the protein over wild type.

Of course, this conferred improvement in stability is predicated on the accuracy of the mAF-min method. To help validate the results, we run molecular dynamics on En-HD variants, which has been done extensively to study the stability of this particular protein.[47,48] Such computational studies have observed several thermal equilibrium ensembles on shorter timescales sampled than this paper.[49] In order to effectively sample the range of unfolding pathways and resolve differences of 10°C in melting temperatures, a large ensemble of replicas with randomized starting velocities is needed. Five replicates of 1 microsecond dynamics were run at a number of temperatures for each of the wild type protein, the NOMELT optimized variant, and a previously engineered variant to >98 °C.[47] The RMSD over each simulation was averaged, yielding a distribution of 5 time-averaged values per protein per temperature. Note that the starting structure (prior to equilibration) for the NOMELT variant is an AF2 predicted structure. In Figure 6, the average RMSD relative to 298K is given as a function of temperature. We can see that the literature engineered variant, “UVF,” retains a tight distribution of displacement around its starting structure up to 370K, while the wild type protein begins to open up at its melting temperature of 56 °C, indicated by a tight distribution of RMSD values widening and an increase in magnitude discontinuously. The NOMELT variant retains its structure until at least 66 °C before doing the same.

**Figure 6:**
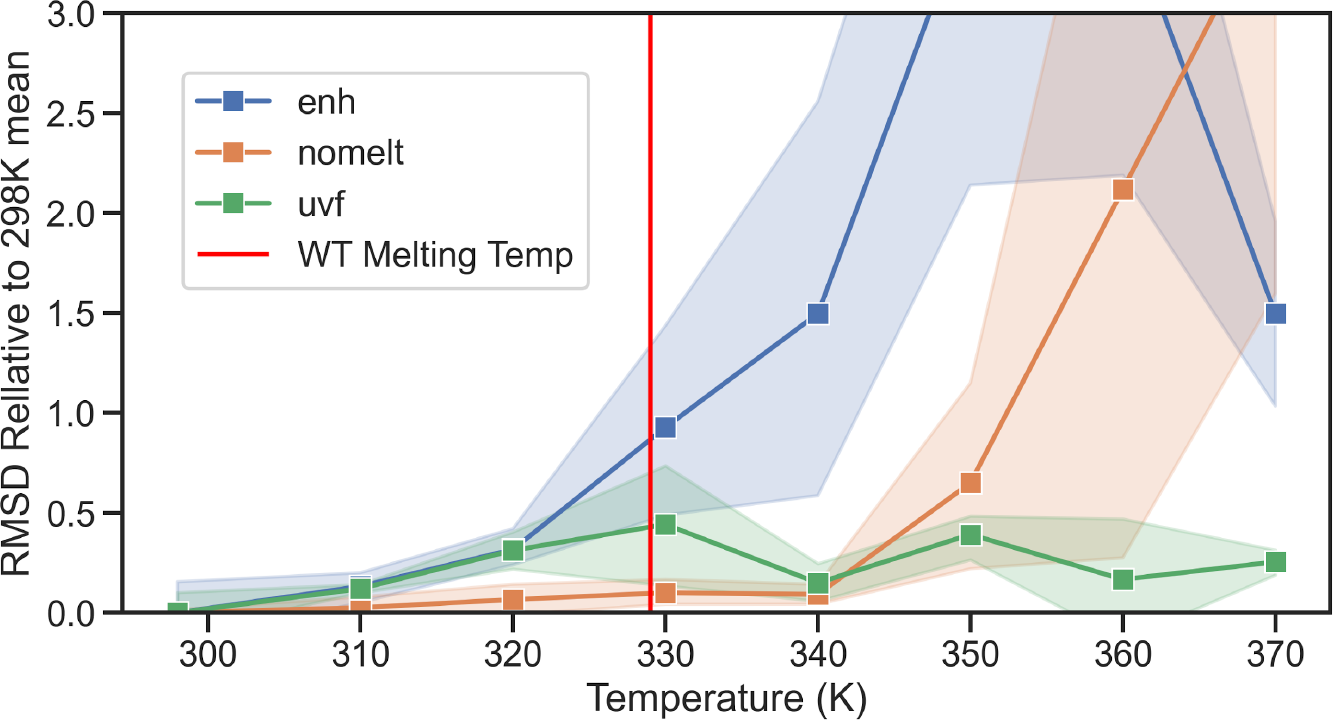
RMSD over 1 microsecond dynamics simulations of En-HD variants. RMSD values are relative to the average value of the RMSD of the variant over the 296K simulations. Distributions are over 5 independent simulations. The UVF variant, experimentally determined to have >98 °C melting temperature, remains tightly bound to its initial structure at all temperatures. The wild type protein opens up from its starting structure at its melting temperature, while the NOMELT variant retains its structure at least another 10K.

#### Case Study 2: NOMELT as a zero-shot estimator of thermophilicity

The output token likelihood of the decoder is a way to estimate the “thermophilicity” of a target mutation or variant according to the model’s learned distribution. Strictly speaking, the model is not an estimator of high temperature stability, but we consider the possibility that in order to recapitulate thermophilic sequences, the model must have some intuition about what makes a protein stable at high temperatures. The idea of using protein language models either zero-shot[22,50] or after self supervised training on a protein family[21,51] to evaluate fitness even when labeled fitness data is unavailable is not novel. However, unlike the family specific techniques, NOMELT is pretrained and does not require an MSA input, and unlike both methods, NOMELT is trained to specifically consider high temperature proteins. For example, ESM2 or MSATransformer are trained on all available proteins sequences and model protein diversity across the evolutionary tree.[24,52] Models like these can be helpful to evaluate fitness by filtering out variation that is universally unnatural, but we argue does not account for interactions that may be specifically important at high temperatures.

Here, we take protein variants with experimental data and evaluate the relationship between NOMELT likelihoods and ground truth stability data. The parent or wild type protein is given to the encoder, and a variant given to the decoder, which predicts a probability vector. After aggregating, the quantity is normalized by the wild type value. See Methods for mathematical details. Firstly, the model was used to rank variants of LovD and LipA variants for which experimental melting temperature was measured, with up to 29 and 12 mutations away from wild type respectively.[53,54] We observe correlation with the experimental melting temperature, as shown in Figure 7. Note that these variants have a minimum E-value of 1.0 with respect to the training set, so are novel to the model. For LovD, NOMELT is extremely predictive out of the box, with a pearson correlation of 0.94 to the true melting temperature. For LipA, predictions are poorer; the model can qualitatively rank a few high temperature variants, but it overestimates the wild type. We hypothesize that this is because the starting melting temperature of the LipA protein is already quite high, and the model was trained on protein pairs where growth temperature was not continuously differentiated, E.g. the model was not told the difference between a 65 °C protein and a 80 °C one. This suggests that NOMELT has some ability out of the box to rank protein variants in terms of high temperature stability, or more accurately, thermophilicity.

**Figure 7:**
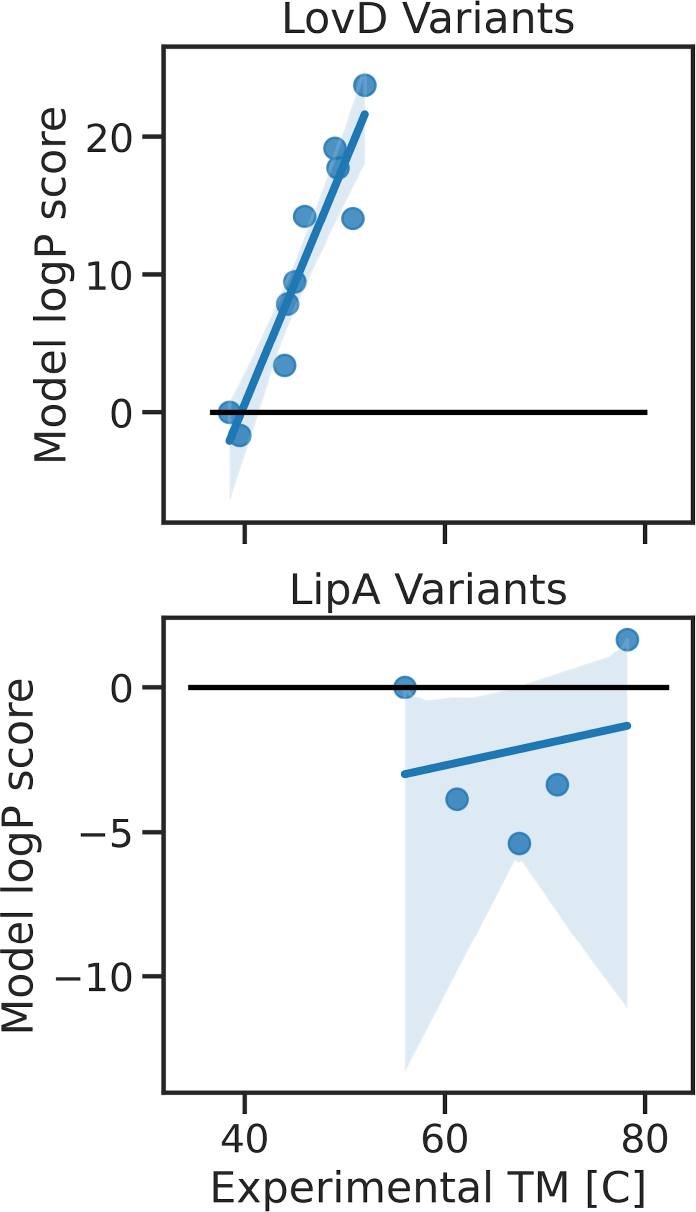
Experimental melting temperature measurements vs NOMELT logP sequence scores for two protein variants sets. Black line is the wild type score. A) NOMELT is highly predictive of LovD variants, with a low wild type melting temperature. B) For LipA variants, the model can qualitatively rank some of the high temperature variants, but overall ranks the wild type sequence, which is already a relatively high melting temperature, too high.

Lipase has also been the subject of a deep mutational scan with catalytic half activation temperature as a target.[55] This experiment of 2,172 single point mutants serves as the only benchmark in ProteinGym with a thermal stability target.[22] When the model is used to rank variants, we note a Spearman correlation to the true target of 0.32, which is state of the art for models that do not require MSAs or homologs as inputs. NOMELT matches the score of ESM2, which was trained on significantly more data, indicating that focusing on thermophilicity has merit.[22] Note that in the original data, replicates were taken for each variant, while ProteinGym only considers the mean. When we consider only the 34% (N=1.7 mil) of pairwise comparisons with T-test P-value <0.05, NOMELT qualitatively distinguishes the proteins 64% of the time. This is impressive in that variants in the DMS library typically have less than 2°C differences.

## Methods

### Dataset

We leveraged our recent dataset, learn2thermDB, to train and test this model.[56] The data contains pairs of homologous proteins across mesophilic and thermophilic temperature regimes. As a first pass, we rigorously filtered this data to contain the strongest possible protein pairs. We only considered pairs with a >20°C difference in optimal growth temperature, with the thermophilic source organism of at least 60°C and mesophilic less than 40°C. The protein pairs themselves were filtered such that only those with >95% sequence alignment coverage of both strands were kept, and the difference in sequence length less than or equal to 10%. This produced 4.3 million protein pairs.

The dataset was clustered using MMSeqs2 to 50% identity on the mesophilic sequences. The parameters used were as follows: min-seq-id=0.5, cluster-reassign=1, cluster-steps=5, s=7, max-seqs=1000, c=0.95, cov-mode=0, similarity-type=2, e=1e-3, cluster-mode=1.[57]

Train, validation, and test sets (80/10/10) were split by cluster such that no sequences in a particular data fold are within similarity thresholds of any sequence in another data fold. For the test set, we randomly sampled 1000 clusters and selected from each a single random sequence. This reduces the bias of large clusters on the evaluation.

### NOMELT Model

NOMELT is an encoder-decoder style transformer neural network. We started from the pretrained ProtT5 foundational protein Language Model, trained to mask-fill protein sequences on the BFD database.[58] The HuggingFace ecosystem of code was used to write the training scripts.[59] Out of the box, the model can be used to generate latent embeddings of proteins, and using a causal language modeling head, the decoder can recapitulate an encoded protein sequence with >99% accuracy. By starting from this model, we leverage general protein knowledge that has been shown to be captured by large protein language models, and we save a significant amount of model training carbon cost. In this work, the model was fine tuned to instead causally generate a thermophilic protein homolog of an input mesophilic protein.

We trained the model in a supervised sequence-to-sequence manner using categorical cross entropy loss on each amino acid in the output sequence in a causal, teacher-forcing strategy. We used a linear ramp-up, ramp-down learning rate and an Adam optimizer with maximum learning rate 1e^-4^ and a ramp rate of 10% of 3 epochs. The model was stopped according to early stopping on validation set loss using a patience of 4 and an improvement threshold of 0.1. This required 7 NVIDIA a40 GPUs 6 hours before the validation loss stopped improving, and cost approximately 2.4kg of carbon emissions. DeepSpeed ZeRO Stage 3 was used to shard and parallelize batches, parameters, and optimization states. BF16 parameter precision was used. A batch size of 140 per device with 2 accumulation steps, was used, for an effective batch size of 1,960.

### Model likelihood

We refer to the softmax probability distribution over amino acids output by the causal model at a specific position *i* in the sequence *P* _*i*_ (• |*x*_0_, *x*_1_, …, *x* _*i*−1_) as *P*_*i*_ where *x*_*i*_ is the amino acid identity at position *i*. The normalized probability of a particular amino acid *j* from the vector at that position is written as *P*_*i*_ (*j*). Following recent work that leverages the probability of language models to evaluate the fitness/target quality due to mutation or of a particular sequence,[21,60] we measure the log likelihood of a sequence or a set of mutations at specific positions according to equation 1 below, where *M* is some set of positions in the sequence:

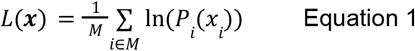

Thus, we can measure the likelihood of a specific mutation at a particular position given a fixed set of all previous amino acids, the combined effect of a set of mutations, or to evaluate the causal likelihood of an entire sequence of arbitrary length. This quantity is useful when comparing to another mutation set or sequence, such as evaluating the likelihood of a mutation occurring relative to the wild type, WT, in equation 2 below:

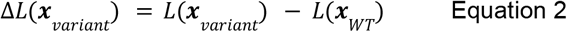

Here, a positive value indicates an increase in likelihood relative to the wild type according to the model. Keeping in mind that the model was trained to generate a thermophilic looking sequence, this quantity carries a different meaning than seen before in canonical zero-shot predictors. This quantity does not represent the estimated likelihood of the mutation/variant among variation, but instead the estimated likelihood of the mutation/variant resembling a thermophilic homolog of the input mesophilic sequence. Note that for evaluating variants that do not have insertions or deletions, we only compare probabilities at mutated residues to the wild type, while instead we must consider the whole sequence probability for instances where there are variable residue counts in variants.

### Model and natural entropy

The categorical cross entropy of the model for some ground truth residue in a sequence at position *i*:

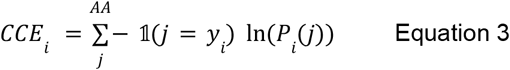

Where AA is all amino acids, plus a number of additional tokens with near 0.0 probability that can be generally discarded. The true amino acid label is *y*_*i*_. This quantity is averaged over all tokens in the dataset and considered the loss function of the model. In order to compare this quantity evaluated on the test set to nature, we took each thermophilic test-set sequence and built an MSA using jackhammer against all other thermophilic sequences >60°C OGT.[33] Search parameters are listed below. For each column in the MSA that appears in the original sequence, we compute a distribution over natural diversity, *P*_*i*_, and use this vector to evaluate the test set entropy on natural variation. This represents a model that treats each position independently instead of causally, and relies upon already existing homologs.

Parameters for the jackhmmer search were F1=0.0005, F2=0.00005, F3=0.0000005, incE=0.0001, E=0.0001, and N=1 (phmmer). We used 32 cores.

### Disulfide bonds

For probing the model’s understanding of Disulfide bonds, we view the test dataset on a teacher-forcing residue basis and consider only positions within the dataset where the true thermophilic sequences have a cysteine. For each of these positions (N=1,687) we measure *P*_*i*_ (*Cystein*) according to the model. The ESMFold predicted structure of each thermophilic test sequence is used to identify which cysteines are likely forming Disulfide bonds. Here, we use the heuristic of 7.5 Å between cysteine C_α_ to label a Disulfide bond.[41] The second cysteine in bond-forming pairs, causally, are labeled as Bond-forming, while all others are labeled as non-bond forming. Note that the model is only aware of residues to the left in sequence, thus from the model’s perspective, the first cysteine in a bonded pair occurring in the sequence is not forming a bond at the time of inference. We computed the one-sided T-statistic between ln (*P*_*C*−*bond*_ (*C*)) and In (*P*_*C*−*non*−*bond*_ (*C*)).

### Sequence Alignment

BLOSUM62 was used to score all sequence alignments.[36] For searching case study proteins against the training set, BLASTp with the following parameters were used: evalue=2.0, word_size=3, qcov_hsp_perc=80.[45] For aligning model designs to ground truths, full Smith-Waterman was used using BioPython with the following parameters: match_score=1, mismatch_score=-1, gapopen=-4, gapextend=-1, penalize_end_gaps=false.[61]

### Structural prediction

For visualization purposes and Disulfide bond estimation, we use ESMFold.[24] For the mAF-min method, we use AlphaFold using the same parameters originally described e.g. reduced database size and no force field relaxation.[23,42]

### Thermal stability estimation

The mAF-min method was used in order to qualitatively compare the high temperature stability of variants of a protein.[42] This method qualitatively ranks variants accumulating many mutations with high accuracy, for a number of protein families. Here, we use the method exactly as described in the original work, with the following exceptions: an ensemble size of 25 instead of 100 was used as suggested by the authors, which was shown to have little loss in accuracy. Even with this reduction, the expense of calling AF2 many times is high, and we randomly selected 50 test set examples to contribute to the results depicted in Figure 4. Additionally, to account for small numbers insertions and deletions not present in the original work, we normalize the method output which qualitatively represents free energy of the system by the number of residues. To validate this choice, we conducted the method with sequence length normalization on 5 variants of Ribonucleases with sequences differing in length from 96 to 110 and melting temperatures between 41.1 and 53.2 °C.[62] This resulted in a Spearman’s correlation of -0.999 with P-value 1.4e^-24^ to melting temperature. Note that the original authors did not provide open source code, so the method was rewritten according to their manuscript in house and is given in this work’s repository.

### Molecular Dynamics

GROMACS was used to conduct molecular dynamics simulations.[63] All simulations were set up identically with the exception of temperature. The protein was placed in a cubic periodic boundary box with at least 1nm between protein and box edge to eliminate unwanted protein-protein interactions across periodic boundary conditions. System was solvated with TIP3P and charge balanced with ions.[64] Data presented is from 1 microsecond long production runs after 100ps NPT equilibration. The charmm36m force field was used.[65] The full configuration files used, including additional information such as timesteps, constraints, velocity cutoffs, etc. are given in a separate repository.[66] Five replicates were conducted for each protein variant at each temperature. Using MDAnalysis 2.3, RMSD values were aligned to energy-minimized PDB reference structures and calculated over residues 10-52 on all variants to capture only the core residues of the 3-helix bundle.[67,68] The RMSD values averaged over each 1 microsecond simulation make up an ensemble of five time-averaged RMSD values, which are depicted in the Figure 6 seaborn plot.[69] The RMSD of each simulation was normalized to its ambient temperature behavior by subtracting the variant’s mean simulation RMSD at 298K. This emphasizes the difference of variants over temperatures as opposed to differences between the baseline behavior of the variants.

## Data and Code availability

All of the work presented here is open source and available on GitHub at https://github.com/BeckResearchLab/nomelt.[70] This repository contains both the code used to train and analyze the model presented in this work, as well as a wrapper of the methods discussed for downstream model use to create an easy entrypoint for using the model. The resulting trained model parameters are available on Zenodo.[71] The major steps of the project are tracked in DVC and can be reproduced with a single command after installation, dependency set-up, and assuming the availability of GPU and CPU resources. The MD scripts and results are available on GitHub.[66]

We advocate for carbon tracking of all resource intensive algorithms.[72] We estimated the cost of training the model at 2.4 kg of carbon and a single replicated of our MD simulations at 1.2 kg. In total for multiple rounds of training and simulations, we estimated that this work required 395 kg of excess carbon.

A number of other software tools were used to develop this pipeline toolset, including MMseqs2 for dataset splitting, HuggingFace for transformer training, Seaborn for plotting, BLAST and HMMER for alignments, ESMFold, pyDSSP, and FATCAT for structural analysis, Data Version Control for parameterizing the pipeline and tracking data states, CodeCarbon for carbon emissions tracking, Dask for parallelization, AlphaFold and pyrosetta for reimplementing the mAF-min method, GROMACS for molecular dynamics simulations, MDAnalysis for analyzing the trajectories, Optuna for optimizing over mutation combinations, PyMOL used for 3D protein structure visualization.[23,24,34,35,45,57,59,63,67,69,72–78]

## Acknowledgements

This work was funded under NSF Engineering Data Science Institute Grant OAC-1934292. This research project was facilitated and supported by the University of Washington’s Hyak computer cluster and IT team.

The University of Washington acknowledges the Coast Salish peoples of the land where this work was conducted, the land which touches the shared waters of all tribes and bands within the Suquamish, Tulalip, and Muckleshoot nations.

## Author Contributions

All authors contributed to writing the manuscript.

EK designed the machine learning solution, prepared the data, trained the NOMELT model and conducted downstream analysis.

CP ran and analyzed all MD simulations presented.

HNA executed NOMELT on the LipA DMS dataset from ProteinGym.

MZ contributed to designing MD simulations.

DACB conceived of the idea to use NMT in this manner, organized development of the model, and contributed intellectually to its development.

## Additional Information

The authors claim no competing interests.

## Supplementary Information

### Alignment of NOMELT EnHD Variant to Wild Type

**Figure S1:**
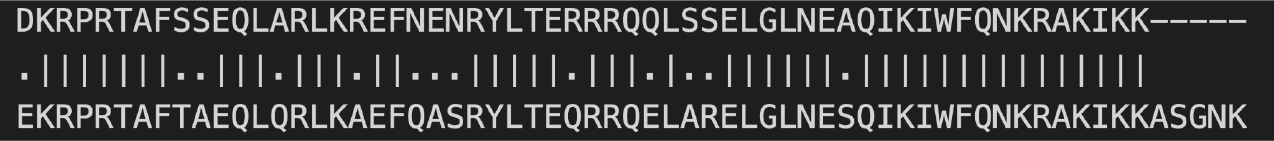
Alignment of wild type EnHD (top) to NOMELT generated variant (bottom).

